# Biocompatible Composite Hydrogel with On-Demand Swelling-Shrinking Properties for 4D Bioprinting

**DOI:** 10.1101/2025.03.26.645296

**Authors:** Peter J. Jensen, Josh P. Graham, Trevor K. Busch, Owen Fitz, Sivani Jayanadh, Thomas E. Pashuck, Tomas Gonzalez-Fernandez

## Abstract

Hydrogels with tunable swelling and shrinking properties are of great interest in biomedical applications, particularly in wound healing, tissue regeneration, and drug delivery. Traditional hydrogels often fail to achieve high swelling without mechanical failure. In contrast, high-swelling hydrogels can absorb large amounts of liquid, expanding their volume by 10-1000 times, due to low crosslink density and the presence of hydrophilic groups. Additionally, some high-swelling hydrogels can also shrink in response to external stimuli, making them promising for applications like on-demand drug delivery and biosensors. An emerging application of high-swelling hydrogels is four-dimensional (4D) printing, where controlled swelling induces structural transformations in a 3D printed construct. However, current hydrogel systems show limited swelling capacity, restricting their ability to undergo significant shape changes. To address these limitations, we developed a high-swelling composite hydrogel, termed SwellMA, by combining gelatin methacryloyl (GelMA) and sodium polyacrylate (SPA). SwellMA exhibits a swelling capacity over 500% of its original area and can increase its original water weight by 100-fold, outperforming existing materials in 4D bioprinting. Furthermore, SwellMA constructs can cyclically swell and shrink on-demand upon changing the ionic strength of the aqueous solution. Additionally, SwellMA demonstrates superior cytocompatibility and cell culture properties than SPA, along with excellent 3D printing fidelity. These findings demonstrate SwellMA’s potential for advanced 4D printing and a broad range of biomedical applications requiring precise and dynamic control over hydrogel swelling and shrinking.

## 1. Introduction

Hydrogels with customizable swelling and shrinking properties have gained significant interest in various biomedical fields. Due to the crucial role of swelling in the absorption of blood and wound exudates, the transfer of nutrients and metabolites, and drug diffusion and release, hydrogels with a high swelling capacity have been extensively used in wound healing^1^, tissue regeneration^2^, and drug delivery^3,4^. However, conventional hydrogels often fail to swell beyond 20% of their original volume without becoming brittle and experiencing mechanical failure. High-swelling hydrogels, defined as materials capable of swelling more than 10-1000 times their initial weight, can absorb large amounts of liquid within their three-dimensional (3D) structure without dissolving, resulting in rapid volume expansion^5,6^. This property is due to the low crosslink density in the hydrogel network and the presence of hydrophilic groups, such as amino, hydroxyl, and carboxyl groups, on the polymer chains^5,6^. Additionally, some high-swelling hydrogels can also shrink in response to environmental stimuli such as pH^7^, temperature^8^, and ionic strength^9^. This stimuli-responsible behavior is promising for different applications such as promoting skin wound closure^10^, promoting cell condensation^11^, triggering drug release^12^, and biosensor design^13^.

The versatility of high-swelling hydrogels has led to their integration into advanced fabrication techniques, such as three-dimensional (3D) bioprinting, enabling the cell-laden constructs for biomedical applications. Utilizing techniques such as extrusion-based bioprinting (EBB), jetting-based bioprinting, and vat photopolymerization, 3D bioprinting replicates biological tissues for applications in tissue engineering, drug delivery, and active cell delivery. EBB uses controlled pressure to extrude hydrogels and cells through a hollow-tip needle, enabling the fabrication of well-defined 3D structures^14^. Jia *et al.* applied this approach to address the need for angiogenesis in living tissues, designing a material that combines GelMA, alginate, and poly(ethylene glycol)-tetra-acrylate using EBB methods^15^. Jetting-based bioprinting builds 3D structures layer by layer using sub-nanoliter droplets of biomaterials and cells, while vat photopolymerization employs ultraviolet (UV) or visible light lasers within a resin bath to generate high-resolution constructs^16^. Although these methods provide precise control over structure and resolution, the static nature of printed constructs limits their adaptability to environmental changes *in vivo*^17^.

To overcome this limitation, an emerging approach known as four-dimensional (4D) printing incorporates the dynamic properties of high-swelling hydrogels to induce structural or functional transformations in 3D-printed structures^18,19^. One of the most studied mechanisms in 4D printing involves shape transformation through hydrogel swelling. This is achieved by immersing the 3D-printed constructs in a swelling medium, such as water, phosphate-buffered saline (PBS), or cell culture medium, depending on the specific application^19^. A common approach is to use a two-layer strategy, where hydrogels with varying swelling capacities are patterned in a bilayer configuration^20–23^. This anisotropic swelling induces internal strain, causing the structure to bend towards the side with lower swelling capacity^20–23^. Ding *et al.* explored this strategy by patterning UV-curable hydrogels with varying swelling capacities^23^. These hydrogels were produced by tuning crosslinking parameters such as the UV exposure time, photoinitiator concentration, polymer concentration and layer thickness, allowing precise control over the bending angle after hydration^23^. Similarly, Diaz-Payno *et al.* 3D-printed tyramine-functionalized hyaluronan (HAT, high-swelling) and alginate with HAT (AHAT, low-swelling) bioinks in a bilayer configuration, enabling fine control over the bending curvature of the constructs post-hydration^20^. While these swelling-based approaches enable high-resolution shape changes, these hydrogels exhibit a limited swelling capacity, typically expanding only 2-3 times their original volume. This moderate swelling can restrict their ability to undergo significant shape changes, especially in in biological environments where external forces are exerted on the hydrogels. To overcome this limitation, Hiendlmeier *et al.* used stereolithography (SLA) to 3D print a super-swelling hydrogel composed of hydroxyethyl-methacrylate (HEMA) and sodium polyacrylate (SPA)^24^. Upon hydration, the hydrophilic sodium polyacrylate residues absorbed water, causing swelling up to 20 times the hydrogel’s original weight and enabling it to resist external forces of up to 100 kPa^24^. Although the authors confirmed that the hydrogel is non-cytotoxic, the synthetic nature of its components limits its bioactivity, making it less suitable for physiological applications. Therefore, there is a need for super-swelling materials for 4D bioprinting that can interact more effectively with biological systems and support dynamic cellular processes.

Gelatin methacryloyl (GelMA) is one of the most used UV-crosslinkable natural polymers in the fields of biomaterials, tissue engineering, and 3D printing due to its biocompatibility and favorable rheological properties. GelMA is synthesized by reacting gelatin with methacrylic anhydride, substituting amine and hydroxyl groups on the side chains of gelatin with methacryloyl groups^25^. Gelatin contributes to the cytocompatibility of GelMA by providing abundant integrin-binding motifs and matrix-metalloproteinase-sensitive groups^25–27^. The methacrylation process allows for UV crosslinking of the GelMA chains in the presence of a photoinitatior such as lithium phenyl-2,4,6-trimethyl-benzoyl phosphinate (LAP) and 2-hydroxy-4′-(2-hydroxyethyl)-2-methylpropiophenone (irgacure-2959). Additionally, by tuning its concentration and environmental temperature, GelMA can be used as bioink, offering viscosity and shear thinning properties suitable for extrusion-based 3D printing^28^. Due to these properties, GelMA has been explored for 4D printing applications. Yang *et al.* achieved differential GelMA swelling by adjusting the crosslinking time of 20% GelMA, which allowed for non-reversable folding of 3D-printed films along the z-axis^29^. Similarly, Gugulothu et al. developed a differential crosslinking approach using a photoabsorber to decrease the crosslinking density of GelMA by attenuating UV light exposure^30^. This technique generated structural anisotropy in the 3D-printed constructs, leading to rapid shape deformation upon hydration^30^. However, these techniques, which rely on modulating GelMA crosslinking density, are limited by the low swelling capacity of GelMA and the potential cytotoxic effects of photoabsorbers and prolonged UV exposure. Furthermore, the swelling achieved through these strategies is not reversible, limiting their use in stimuli-responsive applications.

In this study, we aim to overcome these limitations by producing a biocompatible and stimuli-responsive high-swelling composite hydrogel by combining GelMA, sodium polyacrylate (SPA) and poly(ethylene glycol) diacrylate (PEGDA). This high-swelling hydrogel, hereafter referred to as SwellMA, exhibited a swelling capacity over 500% its original area and was able to increase 100 times its initial water weight, significantly exceeding the capacities of other materials used in 4D bioprinting^20,22,23^. SwellMA addresses these issues, exhibiting similar cytocompatibility and cell adhesion to GelMA, along with superior cell culture properties than SPA. Furthermore, SwellMA offered excellent 3D printing properties, enabling the fabrication of constructs with high fidelity that could swell and shrink on-demand while retaining their shape by varying the ionic strength of the aqueous medium. These results demonstrate the potential of SwellMA for a wide range of biomedical applications in 4D printing and beyond, where controlled swelling and shrinking are essential.

## 2 Materials and methods

### 2.1 GelMA synthesis

GelMA was synthesized using a sequential method as previously described^31^. Briefly, 0.25 M carbonate-bicarbonate buffer was produced with 100 mL deionized (DI) water, 2.65 g sodium carbonate, and 2.1 g sodium bicarbonate (Sigma-Aldrich). The solution was heated at 50°C under steering of 50 rpm and its pH was adjusted using 3 M hydrochloric acid (Sigma-Aldrich) and 1 M sodium hydroxide (Thermo Fisher). The solution was brought up to a pH of 9, and then 10 g of 300 g bloom strength, type A gelatin (Sigma-Aldrich) was added. After gelatin dissolution, 1 mL of methacrylic anhydride (Sigma-Aldrich) was added to the mix in equal amounts over 3 h. The solution, after methacrylic anhydride addition, was transferred into 12-14 kDA dialysis tubes, and placed at 37°C for seven days in DI water, which was changed daily. Finally, the solution was transferred to 50 mL centrifuge tubes, frozen at -80°C, and then lyophilized. Methacrylation of gelatin was confirmed through nuclear magnetic resonance (NMR) spectroscopy.

### 2.2 SwellMA and sodium polyacrylate synthesis

SwellMA was produced by combining SPA with PEGDA, Omnirad 2100, GelMA, and lithium phenyl-2,4,6-trimethylbenzoylphosphinate (LAP) to form a homogeneous mixture as described in **Fig. 1**. First, SPA was synthesized through the acid-base reaction of acrylic acid (Sigma-Aldrich) with 16.206 M sodium hydroxide (Thermo Fisher) to yield a molar ratio of 37.0% sodium acrylate and 63.0% acrylic acid, which has been optimized as the greatest swelling ratio for sodium polyacrylate (SPA) in previous reports^32^. Briefly, acrylic acid was added to a 5 mL glass vial with a stir bar, then placed on a hotplate at 50°C with 150 rpm. The 16.206 M sodium hydroxide solution was then added dropwise to the mix. For SwellMA formation, poly(ethylene glycol) diacrylate (PEGDA) M_n_ 575 (Sigma-Aldrich) was added to the SPA at a concentration of 0.25% w/v as a cross linker between SPA chains. Additionally, the photoinitiators Omnirad 2100 (iGM Resins) at a concentration of 1.0% w/v, and lithium phenyl-2,4,6-trimethylbenzoylphosphinate (LAP) (Sigma-Aldrich) 0.5% w/v were added to the solution. PEGDA M_n_ 575 at 0.25% w/v and Omnirad 2100 at 1% w/v were used as crosslinker and photoinitiator respectively based on previous reports^24^. Previously synthesized GelMA was then added to the solution at a concentration of 10% w/v and dissolved for 30 min at 50°C with 150 rpm. For the formation of geometrically defined constructs, GelMA, SwellMA, and SPA solutions were then casted into a mold and exposed to UV light of 405 nm at an intensity of 25 mW/cm^2^ to polymerize the SPA and crosslink GelMA. SwellMA was also crosslinked through UV light exposure at 365 nm with a dose of 12 mJ/mm^2^.

**Fig. 1.**
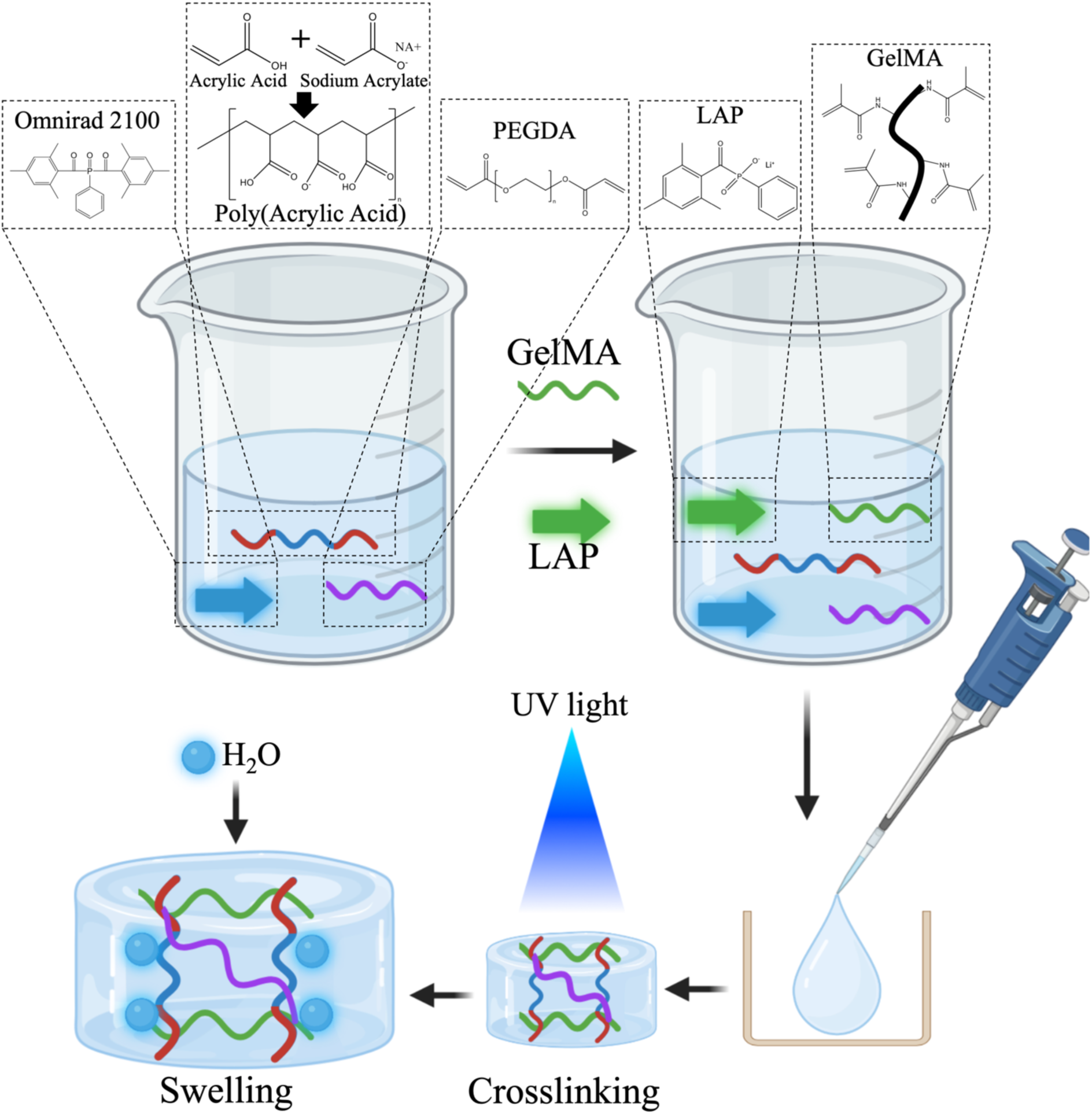
Schematic of SwelMA Synthesis. Poly(acrylic acid) was mixed with poly(ethylene glycol) diacrylate (PEGDA) and Omnirad 2100. Then GelMA and LAP were added to the solution and mixed until a homogeneous mixture was obtained. This solution was pipetted into geometrically defined molds and crosslinked using UV light. After photopolymerization, the resulting constructs were hydrated, causing water absorption and subsequent swelling.

The solution is an aqueous mixture of acrylic acid and sodium acrylate. The SPA solution contains water in a molar ratio equivalent to sodium acrylate, resulting as a byproduct of the acid-base reaction. GelMA is incorporated into this SPA-based solution at a 10% w/v ratio to produce SwellMA. **Fig. 2** provides an example of the molar composition for a 1 mL solution, highlighting that water constitutes only 7.28% w/v. This synthesis does not produce a concentrated SPA polymer but rather an SPA solution, to which GelMA is added for SwellMA synthesis.

**Fig. 2.**
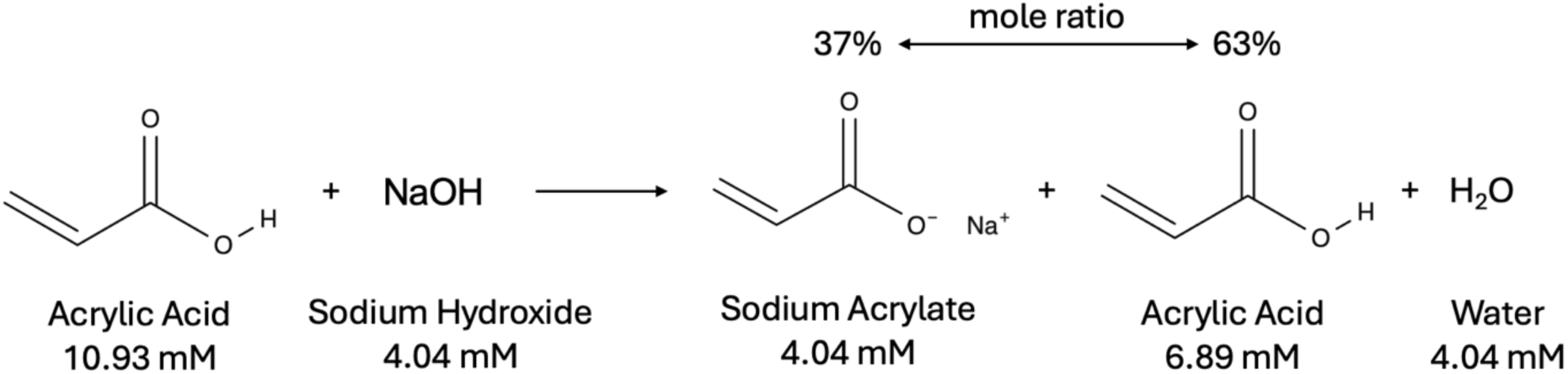
Sodium polyacrylate synthesis. The acid base reaction of acrylic acid and sodium hydroxide yields a mole ration of 37% sodium acrylate to 63% acrylic acid, including a byproduct of water.

### 2.3 Swelling ratio analysis

Samples (n = 3) of 6 mm diameter and 1.59 mm height were casted in a silicone mold with a UV light of 405 nm at an intensity of 25 mW/cm^2^ for 15 s for GelMA, SwellMA, and SPA. The samples were then swelled in 8 mL Dulbecco’s Phosphate Buffer Saline (PBS) (Thermo Fisher) for a 12 h period in a 6 well plate. Each hour the area of a sample was calculated through image analysis. The area of each sample was standardized to their initial areas through Equation 1.

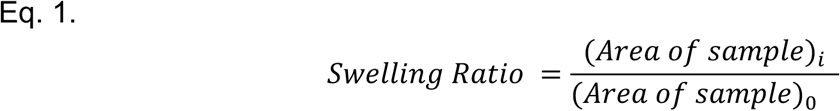

Where *i* is the current area time point and *0* is the initial area of the sample at time 0 h.

### 2.4 Mass swelling ratio characterization

The characterization of the mass swelling ratio was performed through calculating the ratio between water in non-swelled samples *versus* swelled samples after 24 h. Samples were synthesized as described in section 2.3. Non-swelled samples after production or samples after 24 h of swelling (n = 3) were weighed, frozen, lyophilized for 48 h. The mass swelling ratio (SR_m_) was calculated as previously reported^22^, then normalized to the mass swelling ratio of the non-swelled 0 h samples.

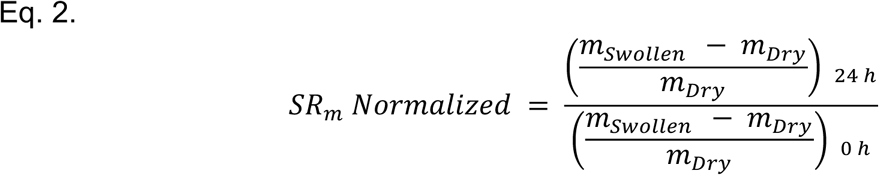

### 2.5 SwellMA swelling and shrinking assessment

Samples (n = 4) were produced as described in section 2.3 and swollen in PBS for 12 h in a 6 well plate. The PBS was then removed, and a 100 mM calcium chloride (CaCl_2_, Sigma-Aldrich, United States) in PBS solution was added to each well, allowing the samples to shrink for 12 h. A timelapse video captured the 12 h swelling and 12 h shrinking cycles. The swelling ratio was calculated at each hourly time point over the 24 h experiment using Equation 1. To optimize the CaCl_2_ concentration, this process was repeated using 0, 5, 10, 25, 50, 75 and 100 mM CaCl_2_. Shrinking percentage was determined by normalizing the final gel area to its original area. For cyclic swelling-shrinking, the same process was repeated with 10 mM CaCl_2_ over four consecutive days.

### 2.6 Rheological properties characterization of strain sweep and frequency sweep

The rheological characterization was conducted with a TA Instrument AR 2000 rheometer. Samples (n = 3) of SwellMA, GelMA, and SPA were synthesized and casted in a mold to produce cylindrical constructs of 8 mm diameter and 1 mm height. The samples were allowed to swell in PBS for 24 h. A frequency sweep was performed followed by a strain sweep sequentially on each sample with an 8 mm radius stainless steel geometry. The frequency sweep was conducted on each sample first with a constant strain of 1.0% and temperature of 37.0°C. Frequencies were taken beginning at 0.1 Hz and increasing logarithmically to 10.0 Hz to measure the storage (G’) and loss modulus (G’’). A strain sweep was then conducted after the frequency sweep at a constant 1 Hz and 37.0°C to measure G’ and G’’ from a strain of 0.1% to 100%. Tangent delta (tanδ) was calculated as the ratio of loss (G’’) to storage (G’) modulus.

### 2.7 Analysis of cell viability and adhesion

Cell adhesion and viability were analyzed through fluorescent imaging and flow cytometry respectively. MSCs were obtained by flushing the femur of Norwegian rats and passaging in expansion media, containing 10% FBS and 1% penn strep on tissue culture treated plastic. Rat MSCs were cultured until passage 6 and collected using 0.25% Trypsin, 0.1% EDTA. 100,000 cells were seeded onto pre-swollen SwellMA, GelMA, and SPA gels of 8mm in diameter and 1.02 mm in height. Cell adhesion and morphology was analyzed 24 and 72 h after seeding through staining with Alexa Fluor™ 488 Phalloidin and 4’,6-diamidino-2-phenylindole, dihydrochloride (DAPI) (Thermo Fisher). Fluorescent imaging was performed using a Keyence BZ-X100 microscope and the BZ-X GFP (Excitation 470/40 nm; Emission 525/50 nm) and BZ-X DAPI (Excitation 360/40 nm; Emission 460/50 nm) filters respectively. Cell viability was assessed using viable cell counts obtained from a Beckman Coulture Cytoflex flow cytometer. At 24 and 72 h after seeding, scaffolds were incubated in 0.25% Trypsin, 0.1% EDTA to remove cells which were centrifuged at 500x g to remove residual trypsin and resuspended in PBS. Flow cytometry was performed with a gain of 14 and 20 for forward scatter (FSC) and side scatter (SSC) respectively. Finally, the resulting data was gated based on FSC-area and SSC-area to remove debris yielding the final cell count.

### 2.8 Characterization of SwellMA biodegradation

The biodegradation of SwellMA was characterized through incubation with collagenase type II, which has been previously used to degrade gelatin and GelMA^33^. GelMA, SwellMA, and SPA samples (n = 6) were produced with 6 mm diameter and 1.59 mm height. 2 units/mL of type II collagenase (Thermo Fisher) in PBS were used to degrade the samples, which corresponds with the concentration of collagenase in initial wound healing^33,34^. Three samples for each group were lyophilized and weighed before degradation to obtain the starting polymer weight. The samples (n = 3) were incubated in an oven at 37°C for 14 days. The collagenase solution was changed every 2 days throughout the experiment. The percent remaining was calculated at the end of the experiment by averaging the first three samples, then dividing each sample by the average.

### 2.9 SwellMA 3D bioprinting characterization

For all the prints in this publication a Regen HU R-200 3D bioprinter was used. The Shaper software (RegenHU, Switzerland) was used for all computer automated designs required for 3D bioprinting. The 3D printing characterization of SwellMA was conducted through filament fusion test, porosity test, and cylinder print test as previously defined^35–39^. The parameters for each of the three tests for both gelMA and SwellMA can be found in **Table 1**. The three tests described below give a characterization of the printability of SwellMA in one, two, and three dimensions. The filament fusion test is a linear test that begins with two lines at 0.25 mm distance and increases by 0.05 mm each consecutive line until reaching the max distance of 1 mm (**Supplementary Fig.1A**)^35,36^. The filament fusion test is used to characterize the spreading of the filament being dispensed from the needle head (**Supplementary Fig.1B**). The filament width was analyzed using imageJ. The SPA filament fusion test was printed using the same conditions as the SwellMA filament fusion test. The filament spreading ratio was calculated through Equation 3.

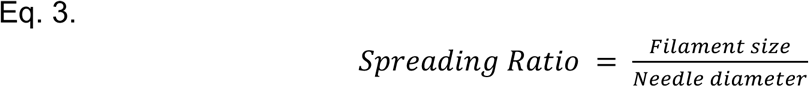

**Table 1.**
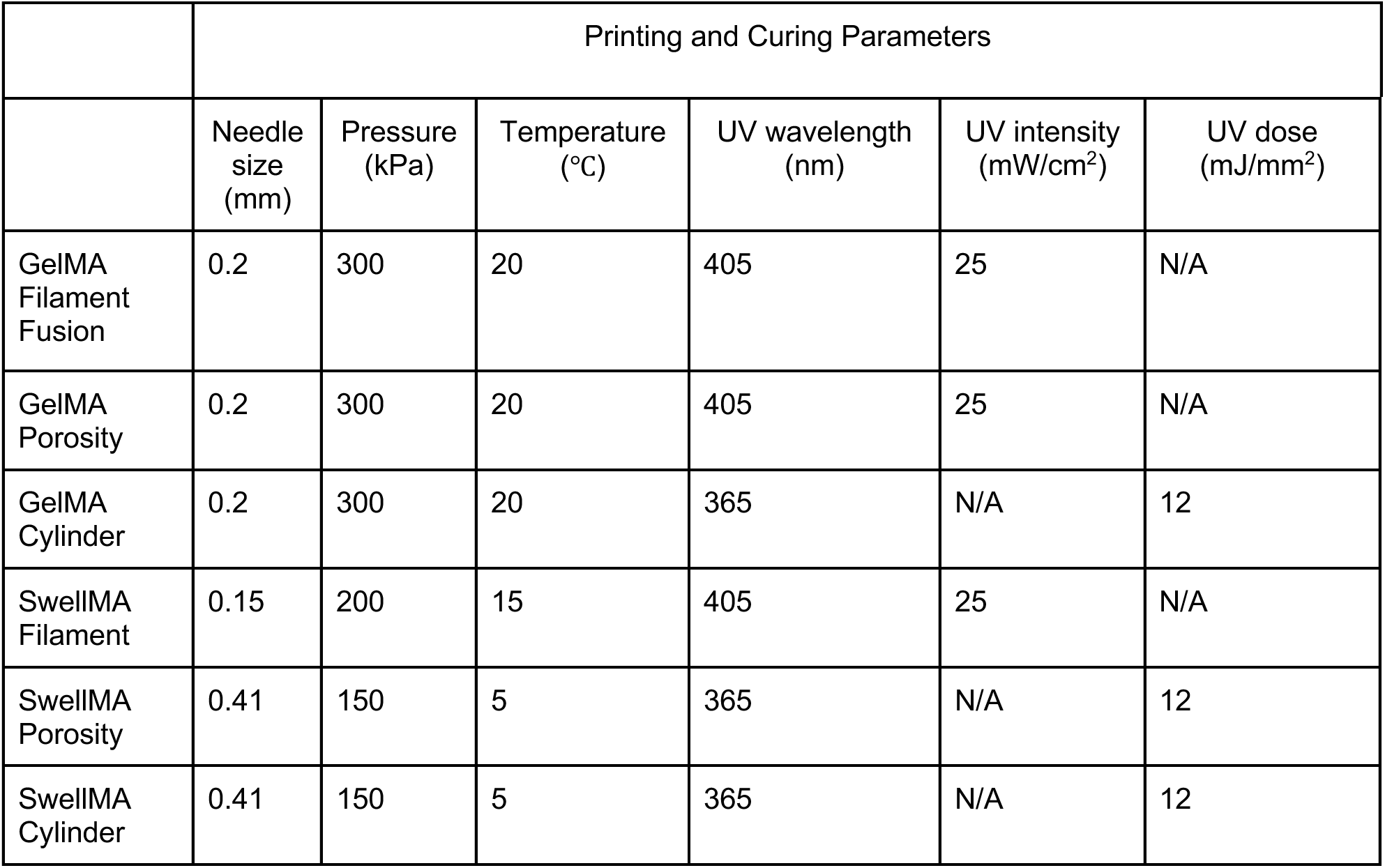
Parameters for printability tests for GelMA and SwellMA.

|  | Printing and Curing Parameters |  |  |  |  |  |
| --- | --- | --- | --- | --- | --- | --- |
|  | Needle size (mm) | Pressure (kPa) | Temperature (°C) | UV wavelength (nm) | UV intensity (mW/cm<sup>2</sup>) | UV dose (mJ/mm<sup>2</sup>) |
| GelMA Filament Fusion | 0.2 | 300 | 20 | 405 | 25 | N/A |
| GelMA Porosity | 0.2 | 300 | 20 | 405 | 25 | N/A |
| GelMA Cylinder | 0.2 | 300 | 20 | 365 | N/A | 12 |
| SwellMA Filament | 0.15 | 200 | 15 | 405 | 25 | N/A |
| SwellMA Porosity | 0.41 | 150 | 5 | 365 | N/A | 12 |
| SwellMA Cylinder | 0.41 | 150 | 5 | 365 | N/A | 12 |

The porosity test is a 20 mm by 20 mm grid with 2 mm by 2 mm squares within (**Supplementary Fig. 2A**). The porosity test provides a two-dimensional analysis of the printability characterization involving the ratio of the experimental pore size to the expected pore size, as illustrated in (**Supplementary Fig. 2B**)^35,36^. The pore area of the print was analyzed using imageJ. The porosity ratio is calculated by Equation 4;

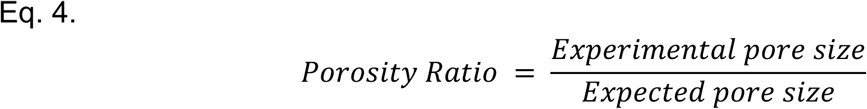

The cylinder print test is a 5 cm diameter cylinder printed to a height of 5 cm (**Supplementary Fig. 3A**)^38,39^. The height of the cylinder was measured using imageJ, which can then be used to calculate the height ratio by Equation 5;

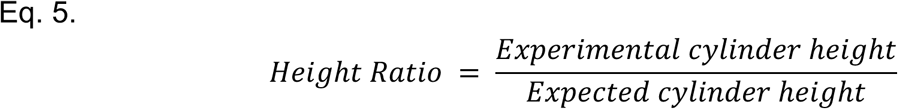

The interior area of the cylinder is measured in the same manner as the porosity test to provide a ratio of the experimental area size as a ratio of the expected pore size (**Supplementary Fig. 3B**).

### 2.10 Statistical Analysis

Data was checked for normality with a Shapiro-Wilks test. Statistical significance was tested with a one-way analysis of variance with a *post hoc* Tukey test for ungrouped data. For grouped data a two-way analysis of variance with a *post hoc* Tukey test was utilized. *P* < 0.05 was considered significant for all statistical tests.

## 3. Results

### 3.1. Characterization of SwellMA swelling and shrinking properties

To maximize the swelling capacity of SwellMA in biological fluids, the swelling ratio was evaluated in PBS at various concentrations of PEGDA crosslinker. Increasing concentrations of PEGDA decreased the swelling ratio and swelling area of SwellMA gels **(Fig. 3A and B)**. A similar effect was also achieved when SPA gels were fabricated with increasing concentrations of PEGDA **(Supplementary Fig. 4)**. SwellMA gels containing 0.25% PEGDA were able achieve a maximum swelling ratio of 5.02 ± 0.15 times their initial area after 12 h of swelling, a 500% increase **(Fig. 3A)**. Additionally, the time required by SwellMA containing 0.25% of PEGDA to achieve maximum swelling was higher than SPA containing the same PEGDA concentration, suggesting that the presence of GelMA decreases SPA swelling capacities **(Fig. 3A and B)**. The comparison of the maximum swelling area in SwellMA and SPA with similar concentrations of PEGDA confirmed the negative effect of GelMA over SPA swelling capacity **(Fig. 3C)**. Despite this observed reduction in maximum swelling capacity in comparison to SPA, SwellMA gels were able to increase 100 times their water weight after hydration in comparison to GelMA, which showed an ineligible increase **(Fig. 3D)**, demonstrating the high-swelling capacity of this material. An additional characteristic of SPA swelling behavior is its capacity to swell and shrink depending on the ionic strength of the surrounding solution. To evaluate the capacity of SwellMA to swell and de-swell, SwellMA, SPA and GelMA hydrogels were swelled for 12 h in PBS and then exposed to a 100 mM CaCl_2_ solution **(Fig. 3E and F)**. After CaCl2 exposure, SwellMA and SPA gels rapidly shrank, recovering their initial area after 3 h **(Fig. 3E and F).**

**Fig. 3.**
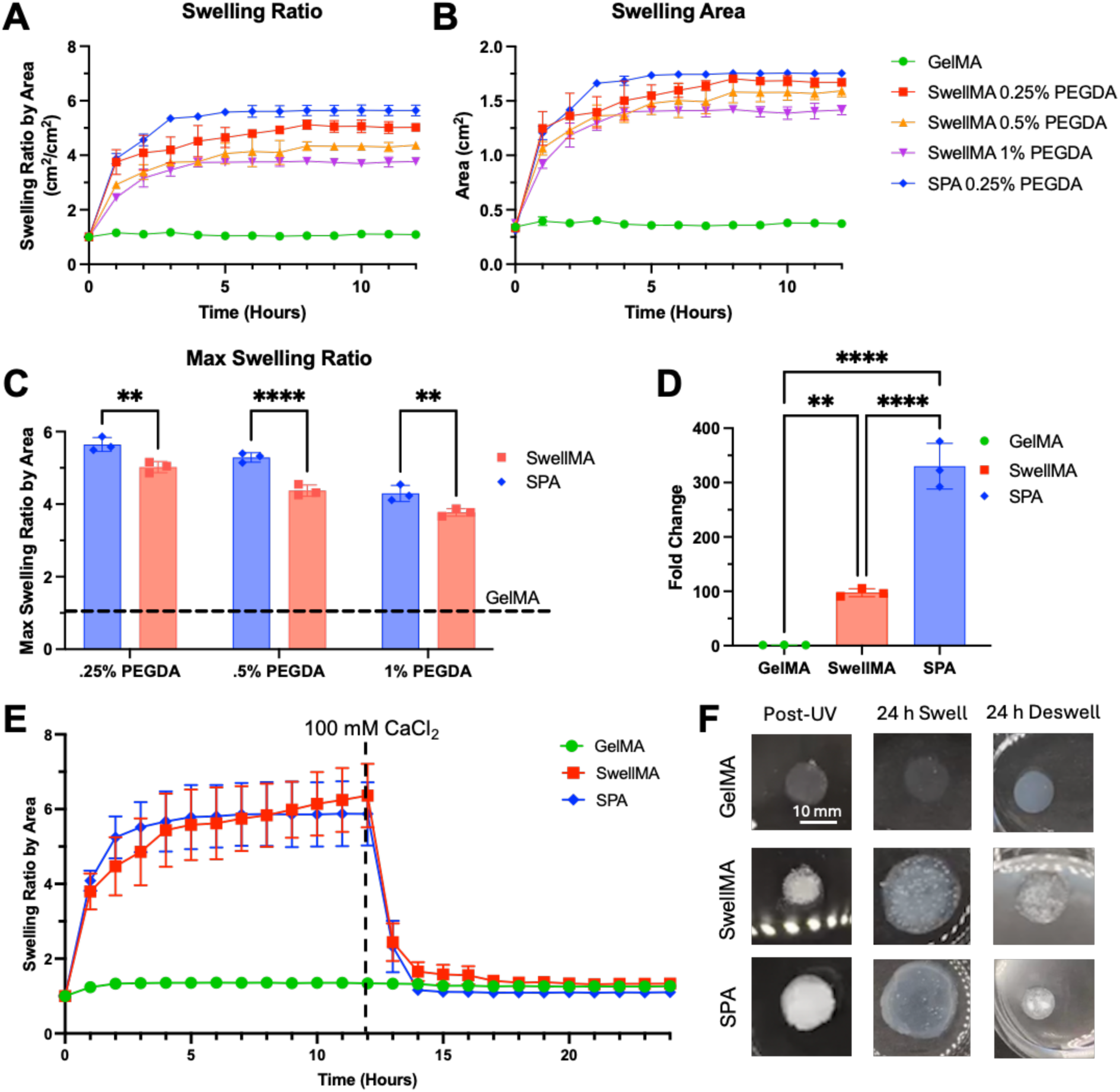
Swelling characterization of SwellMA. **(A-B)** The swelling ratio and area of cylindrical samples swelling over 12 h, **(C)** with a max swelling ratio comparison of SwellMA, SPA, and GelMA. **(D)** The fold change in water weight, normalized to non-swelled day 0 samples. **(E)** Swelling ratio measurements over the course 12 h in PBS, then swelled in calcium chloride for 12 h. **(F)** Representative images of the samples swelling and deswelling over 24 h, scale bar = 10 mm. **** Denotes significance (p < 0.0001), ** (p < 0.01), n = 3 in A-D and n = 4 in E, data shown as mean ± SD.

To determine the minimum CaCl_2_ concentration required to shrink the swollen SwellMA gels, they were first swollen in PBS for 12 h and then incubated in 0, 5, 10, 25, 50, 75, or 100 mM CaCl_2_ solutions for an additional 12 h **(Fig. 4A)**. At 5 mM CaCl_2_, the gels shrank by 38.75 ± 5.69% to their initial area, while 10 mM CaCl_2_ induced a maximum 70.04 ± 2.65% shrinkage, which was not statistically different from the shrinkage observed with the 25, 50, 75, and 100 mM solutions **(Fig. 4A)**. Therefore, 10 mM was determined to be the minimum concentration for maximum SwellMA gel shrinking **(Fig. 4A)**. Additionally, the cyclic swelling-shrinking behavior was assessed for GelMA, SPA, and SwellMA gels over four consecutive cycles using 10 mM CaCl_2_ **(Fig. 4B)**. While SwellMA and SPA did not re-swell to their initial swelling area, both exhibited cyclic swelling and shrinking behavior in response to CaCl_2_ addition **(Fig. 4B)**. In contrast, GelMA did not show cyclic swelling-shrinking behavior and remained relatively unchanged **(Fig. 4B)**.

**Fig. 4.**
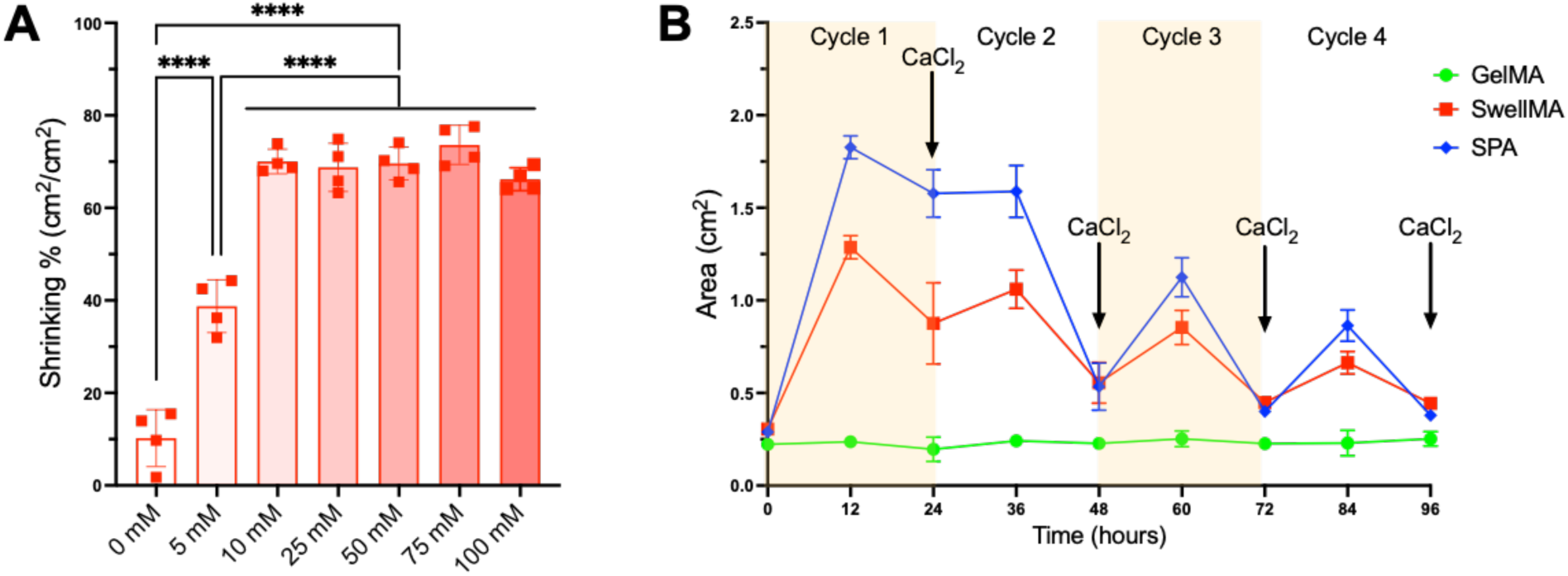
Characterization of SwellMA on-demand swelling and shrinking properties. **(A)** Shrinking percentage of swollen SwellMA gels after incubation in 0, 5, 10, 25, 50, 75, or 100 mM CaCl_2_ solutions for 12 h. **(B)** Cyclic swelling-shrinking behavior of GelMA, SPA, and SwellMA gels over four consecutive cycles, with each cycle consisting of 12 h of swelling followed by 12 h of shrinking through the addition of 10 mM CaCl_2_. ***Denotes significance (p < 0.001), **** (p < 0.0001), n = 4 for A and B, data shown as mean ± SD.

### 3.2. Characterization of SwellMA mechanical properties

After establishing the swelling and shrinking capacities of SwellMA, we assessed the mechanical properties using strain **(Fig. 5A-C)** and frequency sweeps **(Fig. 5D-F)**. The frequency sweeps at a 1 Hz storage modulus for GelMA, SwellMA, and SPA were 1810.33 ± 83.64 Pa, 17833.33 ± 1234.19 Pa, and 3309 ± 237.21 Pa, respectively **(Fig. 5D-F)**. SwellMA consistently showed higher G’ and G’’ values in comparison to GelMA and SPA in both tests. However, SwellMA demonstrated a lower yield point during the strain sweep **(Fig. 5A-C)**. Despite SwellMA transitions to a viscous state earlier GelMA or SPA, its yield point of 21% indicates that SwellMA has suitable mechanics to sustain cell migration, proliferation, and remodeling^40^. Further, the frequency sweep results revealed that SwellMA exhibits a higher tangent delta (tanδ) at 1 Hz **(Supplementary Fig. 5)**, indicating greater viscoelasticity.

**Fig. 5.**
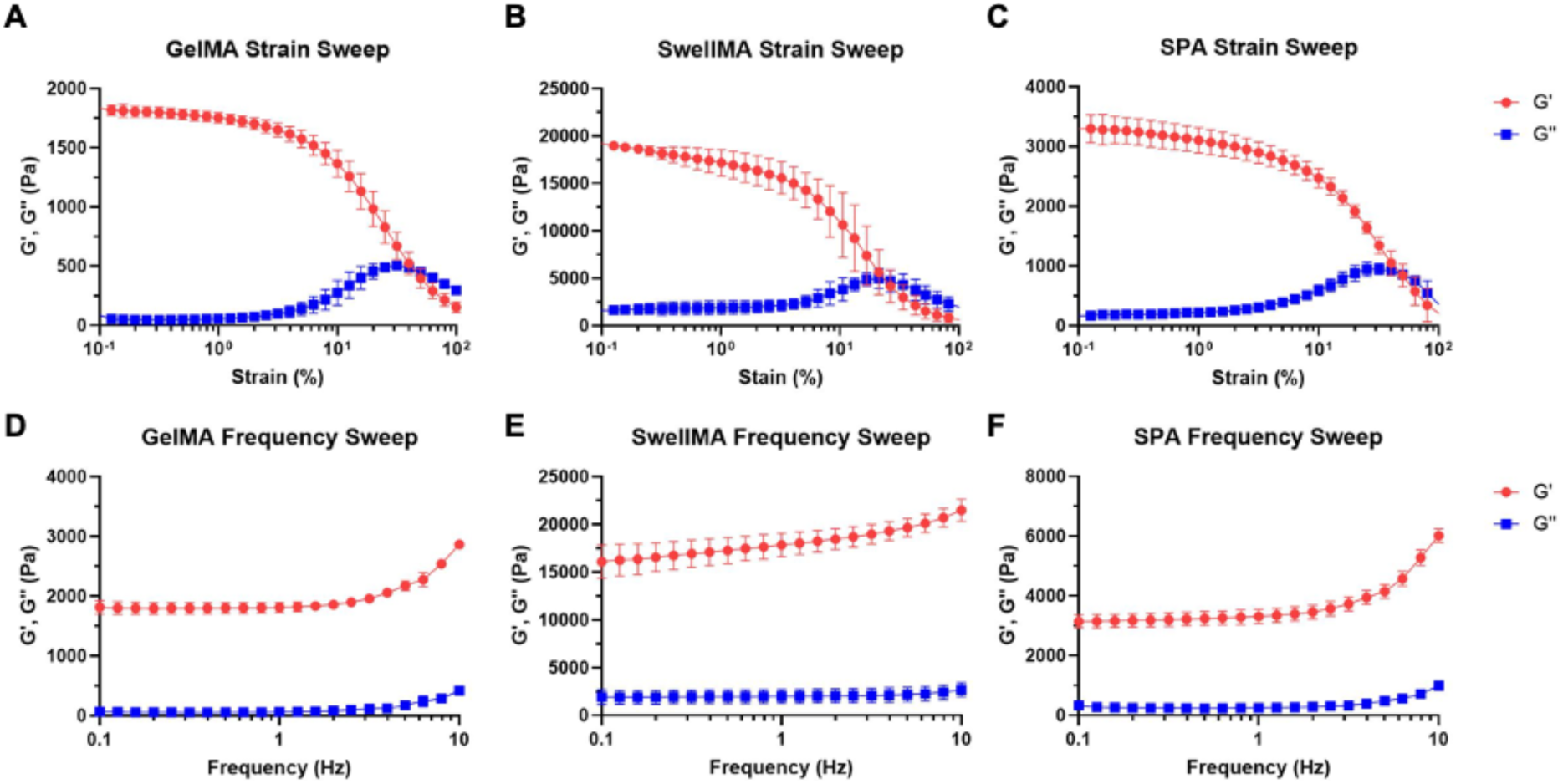
Rheology characterization for GelMA, SwellMA, and SPA. **(A-C)** Strain sweep of GelMA, SwellMA, and SPA with 1 Hz frequency at 37°C. **(D-F)** Frequency sweep of GelMA, SwellMA, and SPA with 1% strain at 37°C. N = 3, data shown as mean ± SD.

### 3.3. Assessment of SwellMA biodegradation and cytocompatibility

GelMA is highly biocompatible due to the presence of integrin-binding motifs for cell adhesion and matrix-metalloproteinase-sensitive groups which allow for its cell-mediated degradation^25^. To test the SwellMA’s biodegradation profile in comparison to GelMA and SPA, cylindrical gels were produced and incubated in collagenase type II for 14 days. Collagenase type II is a matrix metalloprotease (MMP) enzyme that directs collagen remodeling during tissue repair and regeneration^34^. Additionally, collagenase type II has been previously characterized for the degradation of gelatin and GelMA^33^. After 14 days of collagenase type II exposure, the GelMA gels were completely degraded but no differences could be observed in the SwellMA and SPA gels **(Fig. 6A)**.

**Fig. 6.**
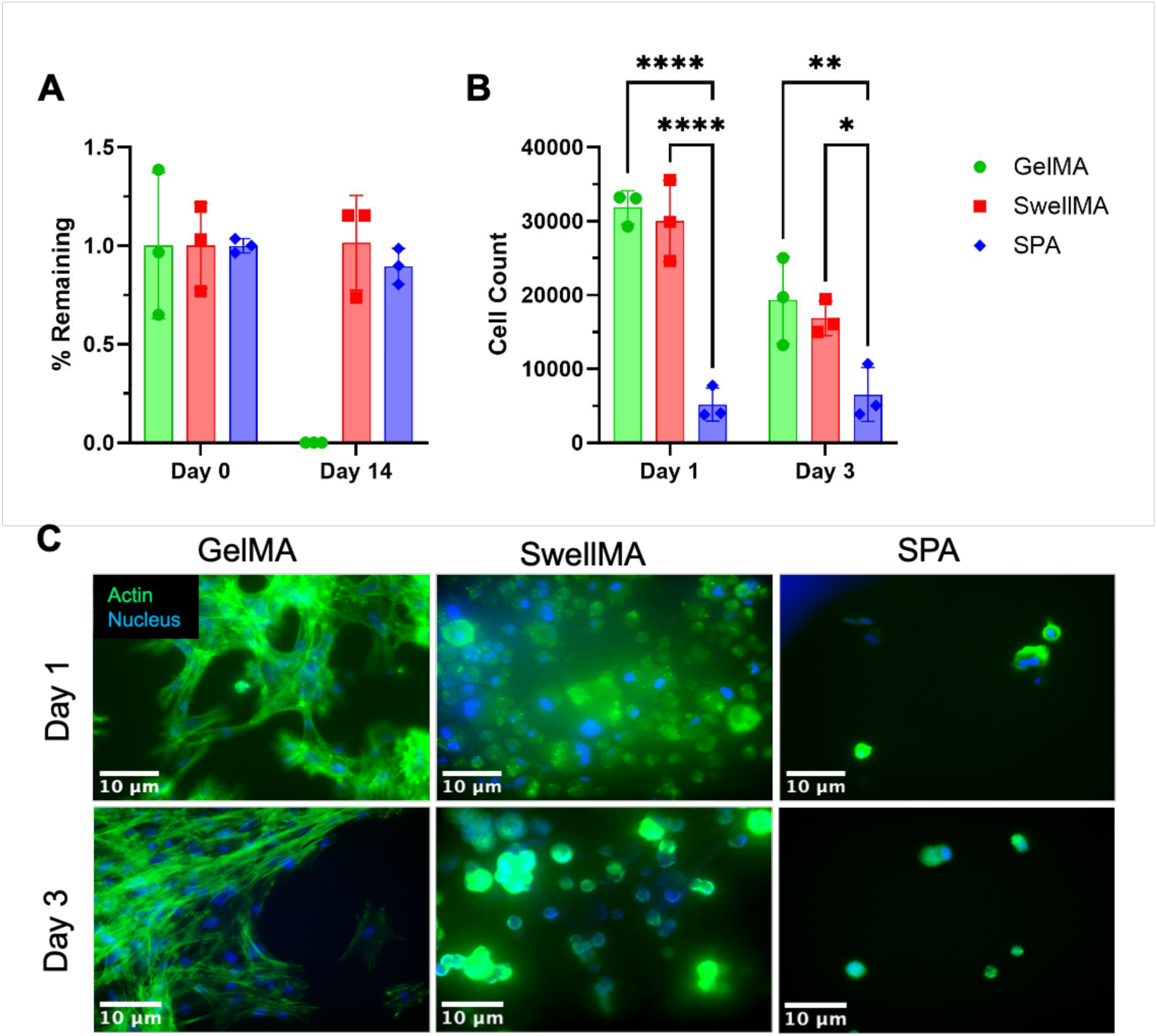
Biodegradation and cytocompatibility analysis of SwellMA constructs. **(A)** Quantification of % of polymer remaining after incubation with collagenase type II over 14 days. **(B-C)** Flow cytometry quantification of cell count and imagining of cells actin and nucleus at days 1 and 3 after seeding (green = actin cytoskeleton, blue = nucleus, scale bar = 10 μm). **** Denotes significance (p < 0.0001), ** (p < 0.01), * (p < 0.05), n = 3, mean ± SD.

In addition to SwellMA’s biodegradation, we also explored SwellMA’s cytocompatibility in comparison to GelMA and SPA. Cylindrical gels were produced and incubated in PBS for 24 h to achieve maximum swelling. After swelling, primary human bone marrow-derived MSCs were seeded on the gels and the cell count and cell morphology was monitored for 3 days **(Fig. 6B and C)**. SwellMA showed similar cell count to GelMA at days 1 and 3 after seeding **(Fig. 6B)**, confirming the cytocompatibility of the material. In comparison, cells failed to adhere and grow in SPA gels due to the lack of cell adhesion domains in this material **(Fig. 6B)**. Cell morphology was also analyzed through staining of the cell cytoskeleton and nucleus **(Fig. 6C).** While cells in GelMA showed a spindle-like morphology with well-defined actin cytoskeleton, MSCs in SwellMA exhibited a round morphology **(Fig. 6C)**.

### 3.4. SwellMA 3D Printability Assessment

GelMA has been widely explored for 3D printing applications due to its tunable rheological properties which allow for high fidelity after extrusion^37,41^. The reversible thermal gelation of GelMA allows for aqueous solutions of 7-10% GelMA concentrations to exhibit viscous shear-thinning properties when 3D-printed at temperatures lower than 20°C^41,42^. To test the printability of SwellMA in comparison to GelMA and SPA, we performed the filament fusion test to characterize the spreading of the filament being dispensed from the needle head^35,36^. SwellMA exhibited high 3D printability in comparison to SPA, which was not printable due to the lack of defined filaments after extrusion **(Fig. 7A)**. Additionally, the filament architecture of 3D-printed SwellMA was preserved after UV crosslinking **(Fig 7A)**. Despite SwellMA’s increased in printability in comparison to SPA, the spreading ratio of SwellMA was significantly higher than GelMA, suggesting lower printing fidelity **(Fig. 7B)**.

**Fig. 7.**
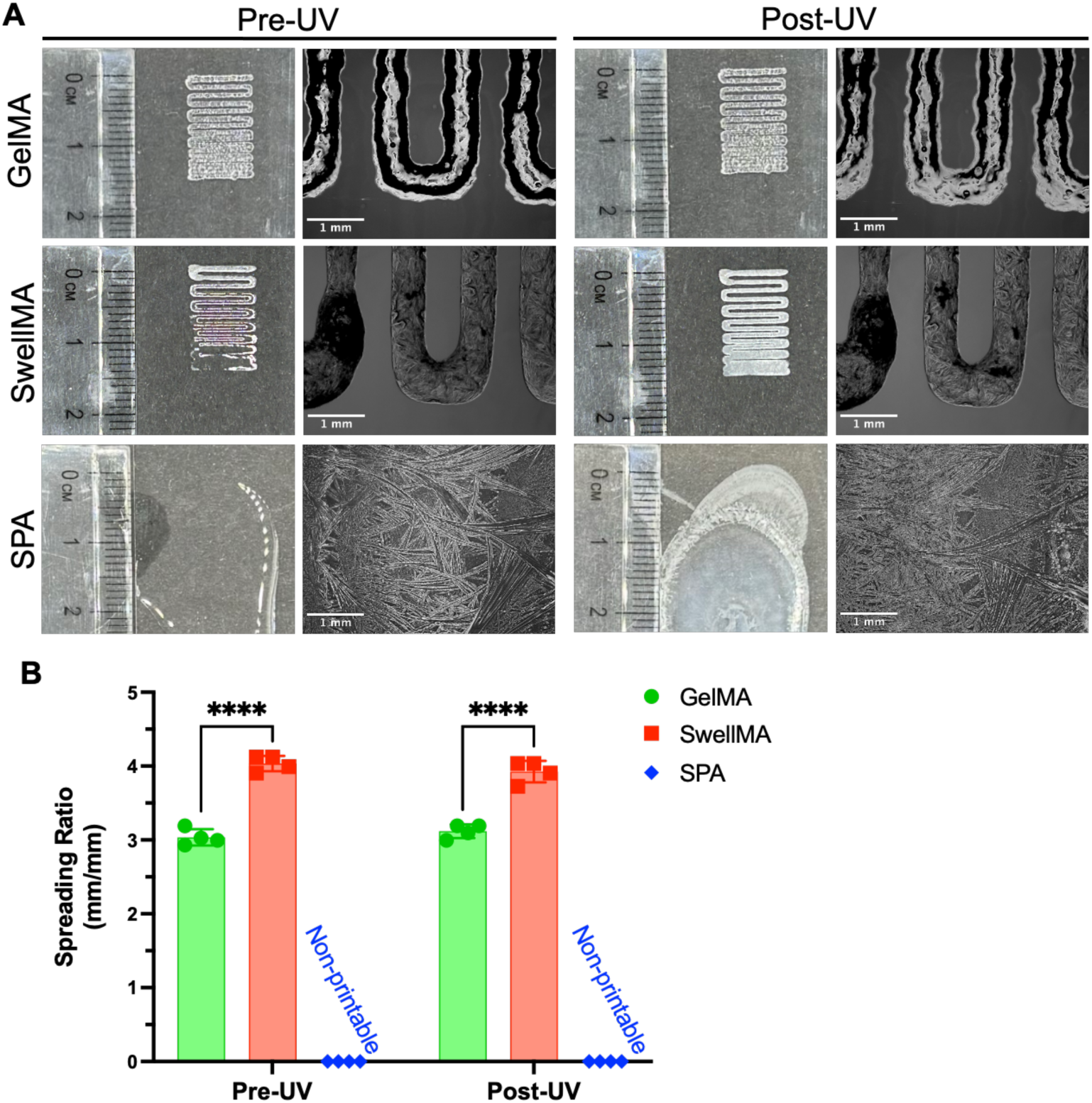
Filament fusion and filament spreading test of GelMA, SwellMA, and SPA. **(A)** Macroscopic and microscopic images of filament before and after UV curing, scale bar = 1 mm. **(B)** Quantification of the filament spreading ratio. **** Denotes significance (p < 0.0001), n = 4, data shown as mean ± SD.

### 3.5. On-demand swelling and shrinking capacity of 3D-printed SwellMA constructs

After characterizing SwellMA’s printability through the filament fusion and spreading ratio tests, we assessed the capacity of SwellMA to 3D print constructs with a defined architecture and the swelling and on-demand shrinking capacities of these constructs. First, we 3D-printed geometrically defined grid constructs with defined internal porosity **(Fig. 8)**. After 3D printing and UV crosslinking, 3D-printed constructs were swelled in PBS for 24 h **(Fig. 8A and Supplementary video 1)**, showing a ∼5 times increase of its initial area (from ∼4 cm^2^ to ∼20 cm^2^), proving that 3D-printed constructs conserve the swelling capacities of casted gels as characterized in section 3.1. To test the shrinking properties of swelled 3D-printed constructs, constructs were incubated in 100 mM CaCl_2_ for 24 h, after which constructs were able to recover their initial shape **(Fig. 8A, and Supplementary video 2)**. This swelling-shrinking capacity was further characterized by calculating the porosity ratio of the 3D-printed constructs **(Fig. 8B)**. After swelling, the porosity ratio of the printed constructs increased ∼5 times its initial value (from 0.37 ± 0.04 to 1.92 ± 0.11) **(Fig. 8B)**. This increased porosity ratio decreased when the constructs were incubated in 100 CaCl_2_ for 24 h, coming back to almost its original value (from 0.37 ± 0.04 to 0.61 ± 0.004) **(Fig. 8B)**. SwellMA’s swelling-shrinking capacity was also compared to 3D-printed GelMA constructs, which did not significantly change their area or porosity ratio after hydration or exposure to 100 mM CaCl2 **(Supp. Fig 6)**. Finally, we assessed the handling and robustness of the swelled 3D-printed constructs, which could be easily picked up with metal tweezers and remained intact after manipulation **(Fig. 8C).**

**Fig. 8.**
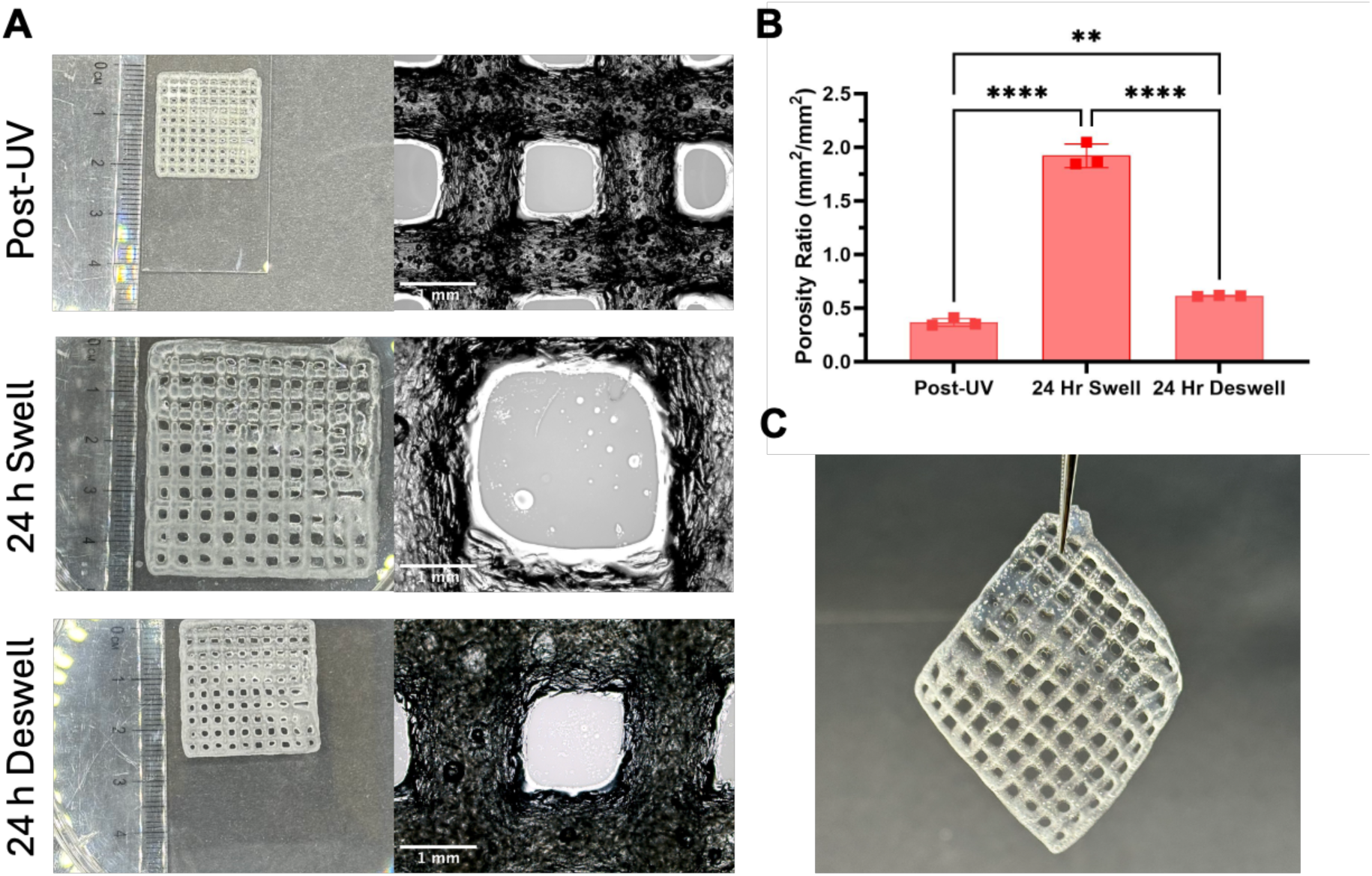
Assessment of SwellMA for the 3D printing of geometrically defined macroporous constructs. **(A)** Macroscopic and microscopic imaging of SwellMA 3D-printed grids after 24 h of swelling and 24 h of shrinking, scale bar = 1mm. **(B)** Quantification of porosity ratio after UV crosslinking, swelling and shrinking. **(C)** Image of 3D-printed SwellMA constructs after swelling when picked up with metal tweezers. **** Denotes significance (p < 0.0001), ** (p < 0.01), n = 3, data shown as mean ± SD.

To further confirm the capacity of SwellMA to 3D print geometrically defined constructs with on-demand swelling-shrinking capabilities, we used SwellMA to 3D print cylindrical constructs and compared their swelling and shrinking properties to GelMA prints **(Fig. 9)**. In comparison to GelMA prints which shape remained unchanged, 3D-printed SwellMA cylinders were able to swell ∼3 times their initial height **(Fig. 9A and B)** and ∼6 times their initial cross-sectional area **(Fig. 9C and D)** after 24h in PBS **(Supplementary video 3)**. Additionally, when the swelled SwellMA constructs were incubated for 24 h in a 100 mM CaCl_2_ solution, their height and cross-sectional area was able to shrink back almost to their initial shape **(Fig 9A-D and Supplementary video 4)**. Therefore, these data demonstrate the superior capacity of 3D-printed SwellMA constructs to swell and shrink on-demand in comparison to GelMA.

**Figure 9.**
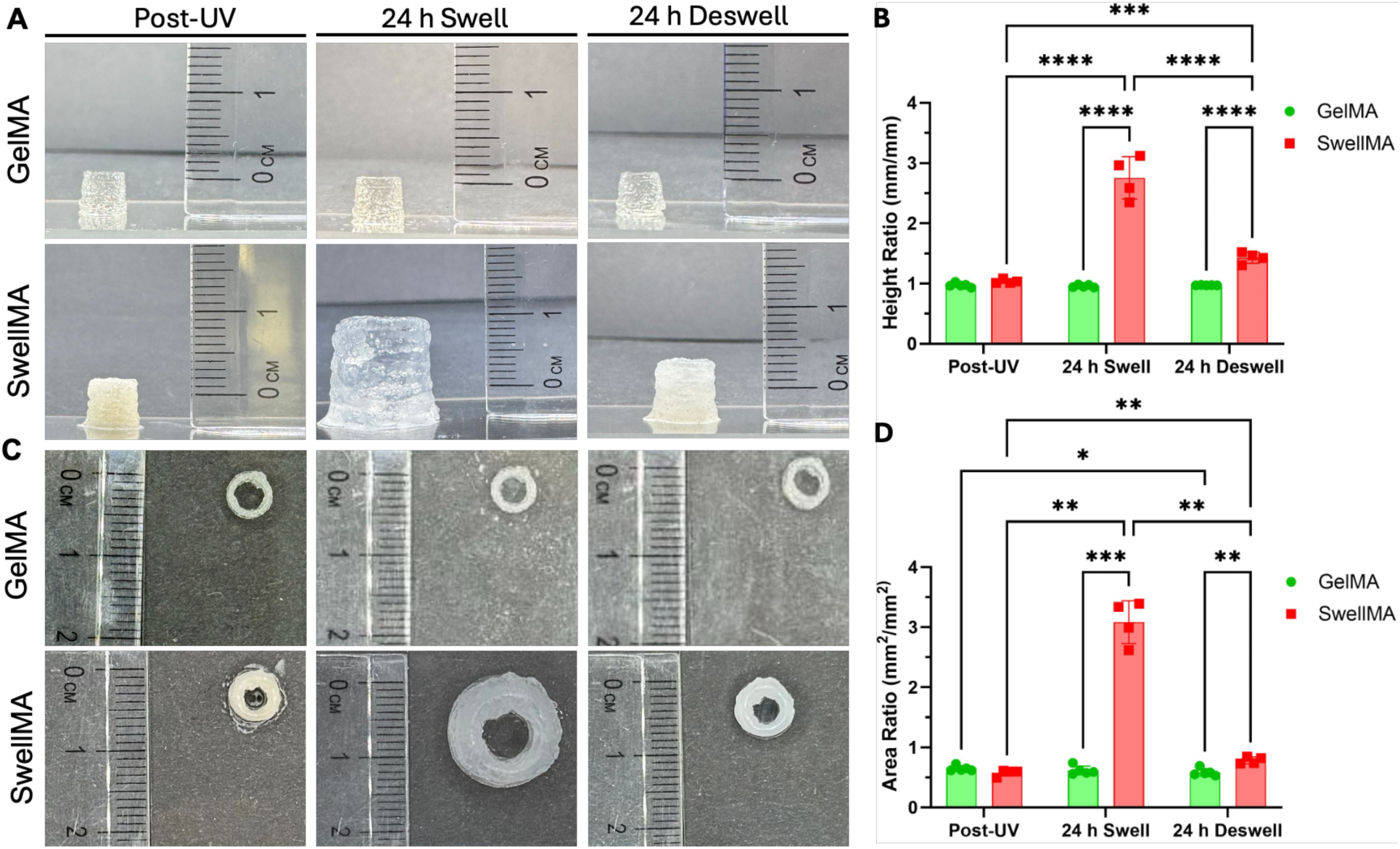
Cylinder print test of GelMA and SwellMA after UV crosslinking and after 24h of swelling and shrinking. **(A)** Macroscopic profile images and **(B)** quantification of cylinder height ratio. **(C)** Macroscopic top view images and **(D)** quantification of the cross-sectional area ratio. **** Denotes significance (p < 0.0001), *** (p < 0.001), ** (p < 0.01), * (p < 0.05), n = 3, data shown as mean ± SD.

## 4. Discussion

In this study, we introduce SwellMA for the first time, a biocompatible, stimuli-responsive, high-swelling composite hydrogel that uniquely combines the super-swelling properties of SPA with the bioactivity and printability of GelMA. SPA was selected for its exceptional ability to absorb 100 to 1000 times its mass in water, due to the presence of sodium and carboxylic groups in its structure^43,44^. GelMA was chosen for its capacity to support cell adhesion and matrix remodeling while providing favorable rheological properties for 3D printing and biofabrication^25,26,41,42^. We synthesized SwellMA by reacting the alkene groups in GelMA, SPA, and PEGDA with photo-initiators, resulting in a material that harnesses the super-swelling properties of SPA alongside the biocompatibility of GelMA. This composite structure is hypothesized based on previous research using UV light to cure a hydrogel made from SPA, PEGDA, and 2-hydroxyethyl methacrylate (HEMA)^24^. However, further structural analysis is needed to confirm the exact bonding mechanism in SwellMA.

SwellMA addresses the need for controlled super-swelling in the field of 4D printing, demonstrating a 500% increase in area, and a 100-fold weight increase. The swelling behavior of SwellMA depended on factors such as salt concentration, the ratio of monovalent to divalent cations in the external solution, the surface area-to-polymer mass ratio, and the polymer crosslinking density^43,44^. Additionally, SwellMA demonstrated on-demand shrinking properties when incubated with CaCl_2_ solutions, with 10 mM CaCl_2_ being the minimum concentration required to achieve maximum shrinking. This concentration falls within a biocompatible range. For example, culture media supplemented with 6 to 10 mM CaCl_2_ has been shown to enhance MSC migration and proliferation^45^. Furthermore, SwellMA exhibited cyclic swelling and shrinking behavior, responding to the addition and removal of CaCl_2_. However, SwellMA was not able to swell back to its initial area after shrinking, which aligns with previously reported limitations of SPA reswelling^46^. Despite this limitation, the dynamic behavior of SwellMA highlights its potential for 4D bioprinting applications and for investigating the cellular effects of dynamic changes in substrate shape and configuration^22,47,48^.

The unique interactions between the components of SwellMA resulted in specific mechanical properties distinct from those of GelMA and SPA. In frequency sweeps, SwellMA exhibited a higher storage modulus than both SPA and GelMA at 1 Hz, along with a yield point of 21% and higher tanδ. The higher storage modulus may result from increased covalent bonding within the polymer network^49,50^. The synthesis method for SPA produces a solution consisting of 37% sodium acrylate and 67% acrylic acid, to which 10% (w/v) GelMA is added. This addition raises the storage modulus from 3309 ± 237.21 Pa in SPA to 17833.33 ± 1234.19 Pa in SwellMA, supporting the hypothesis of increased covalent bonding due to the additional alkene groups in GelMA. The alkene-based network enhances the number of physical bonds capable of storing energy within the gel. The yield point and higher viscoelasticity together suggest that the mechanical properties of SwellMA recapitulate the dynamic viscoelasticity of tissues. For example, chemically modified GelMA with greater viscoelasticity improved cell-material interactions to enhance the chondrogenesis and osteogenesis of MSCs^51^. Furthermore, the storage modulus of SwellMA falls within the range of hydrogels with mechanical properties optimized for the induction of MSC osteogenesis^27^, further highlighting its potential for applications in tissue engineering and regenerative medicine.

To explore the biological implications of SwellMA’s mechanical properties, we investigated its degradation kinetics and assessed its potential as a cell culture platform. SwellMA was incubated in collagenase type II for 14 days and, unlike GelMA, which was fully degraded, SwellMA showed no significant degradation. Previous studies have shown that collagenase-resistant methacrylamide crosslinks in GelMA hinder its degradation compared to collagen hydrogels^33^. The presence of additional methacrylamide crosslinks in SwellMA could explain the material’s reduced degradation. Furthermore, SPA is resistant to enzymatic degradation by collagenases, and it has been reported to inhibit the action of MMP-2, MMP-9 and collagenase during wound healing^52^. SwellMA’s reduced degradation capacity could be advantageous to achieve exudate control in chronic wounds with high proteolytic activity^52^. Cell-SwellMA interactions were also analyzed through human MSC seeding onto SwellMA, SPA and GelMA scaffolds. Cell adhesion was not significantly different between GelMA and SwellMA, but both exhibited greater cell adhesion in comparison to SPA. While the cell numbers on GelMA and SwellMA scaffolds were similar, differences in MSC morphology were observed. Previous studies have demonstrated that arginine-glycine-aspartic acid (RGD)-functionalized substrates with high elasticity, which cannot dissipate cellular forces, inhibit cell adhesion and spreading by preventing ligand clustering^40,53^. Although GelMA is an elastic material due to the covalent methacrylamide crosslinks^54^, its sensitivity to enzymatic degradation allows for cell-mediated matrix remodeling and ligand clustering, resulting in enhanced cell adhesion and spreading^55^. In contrast, the lack of enzymatic degradation of SwellMA, might impair ligand clustering, favoring cell-cell interactions and reducing cell adhesion and spreading. Additionally, the presence of SPA within the SwellMA structure could have increased the distance between cell adhesion domains provided by GelMA, hindering cell spreading and the formation of focal adhesions^56^. Future studies will focus on enhancing cell-SwellMA interactions through adhesion ligand functionalization and by optimizing the proportions of GelMA, SPA, and PEGDA in the mixture.

The favorable swelling, rheological, and cytocompatibility properties of SwellMA were leveraged for additive manufacturing through EBB. Printability was evaluated across all three dimensions through well-established printability tests^36^, using GelMA as the gold standard for comparison. While SwellMA exhibited lower printing fidelity in filament spreading and porosity tests compared to GelMA, its 3D printing performance was superior to SPA, which was non-printable. Future studies will optimize SwellMA’s printing fidelity by adjusting the concentrations of GelMA, PEGDA, and SPA in the mixture and evaluating their impact on its rheological properties. In contrast to 2D printability tests, SwellMA exhibited no significant differences from GelMA in the 3D cylinder printing test. Additionally, the 3D printed SwellMA constructs were able to maintain their shape during swelling and exhibited controlled shrinking in response to variable osmotic conditions, demonstrating its potential for 4D bioprinting. These properties surpass those of previous bioinks with enhanced swelling capabilities. Jia *et al*. explored the use of GelMA, alginate, and PEGTA, a different PEG derivative, for printing defined shapes for angiogenesis applications^15^. However, it lacked the swelling properties necessary for time-dependent morphological changes, requiring the insertion of large 3D-printed structures into biological systems. In contrast, SwellMA offers a minimally invasive implantation alternative. Liu *et al.* addressed the need for minimally invasive technology by developing an amphiphilic dynamic thermoset polyurethane with programmable swelling to restore 3D shapes from 2D patterns^20^. However, cell adhesion was not demonstrated. SwellMA overcomes this limitation by supporting MSC adhesion at levels comparable to GelMA. SwellMA’s unique swelling-shrinking capabilities, rheological properties, and cell adhesion potential position it as a valuable tool for 4D bioprinting. Future studies will explore the 3D printing of GelMA and SwellMA patterns to achieve controlled shape transformations^20–23^. For example, Diaz-Payno *et al*. demonstrated that printing a high-swelling tyramine-functionalized hyaluronan (HAT) bioink as the bottom layer and a low-swelling alginate-HAT bioink as the top layer resulted in an upward concave bending upon immersion^22^. A similar approach could be applied with SwellMA by printing it as the bottom layer for providing high swelling and low swelling GelMA as the top layer to induce a concave curvature transformation.

## 5. Conclusion

In this study we introduce SwellMA, a novel composite material that exhibits significantly higher swelling capacity than GelMA while preserving GelMA’s cytocompatibility and 3D printability. Notably, SwellMA’s on-demand shrinking capability, achieved by adjusting the ionic strength of the surrounding aqueous medium, represents a substantial advancement over previous strategies for enhancing GelMA’s swelling, which lacked the ability for controlled shrinking. This controllable swelling and shrinking feature highlights SwellMA’s potential for 4D bioprinting applications, where shape transformations of 3D-printed structures are triggered by an external stimulus. Future research will focus on enhancing SwellMA’s biocompatibility and printability by optimizing the concentration of its components, and on exploring complex 4D printing transformations through SwellMA-GelMA patterning. Additionally, SwellMA provides a promising platform for studying the effects of cyclic swelling and shrinking on cellular processes, particularly in comparison to other 4D printing materials that do not support on-demand shrinking.

## Acknowledgements

Tomas Gonzalez-Fernandez would like to acknowledge the start-up funds provided by the Department of Bioengineering and the P.C. Rossin College of Engineering & Applied Science at Lehigh University, the Orthoregeneration Network (ON) Kick Starter grant, and the Career Development Award from the American Society of Gene & Cell Therapy. The content is solely the responsibility of the authors and does not necessarily represent the official views of the American Society of Gene & Cell Therapy. Joshua P. Graham would like to acknowledge the support provided through the National Science Foundation Graduate Research Fellowship under Grant No 2234658. TKB would like to acknowledge the funding provided through the IDEAS Experiential Learning Grant and the Forum Research Grant provided by the Lehigh University.

