## Supplementary material for "Biocompatible Composite Hydrogel with On-Demand Swelling-Shrinking Properties for 4D Bioprinting": Supplementary_Information.pdf

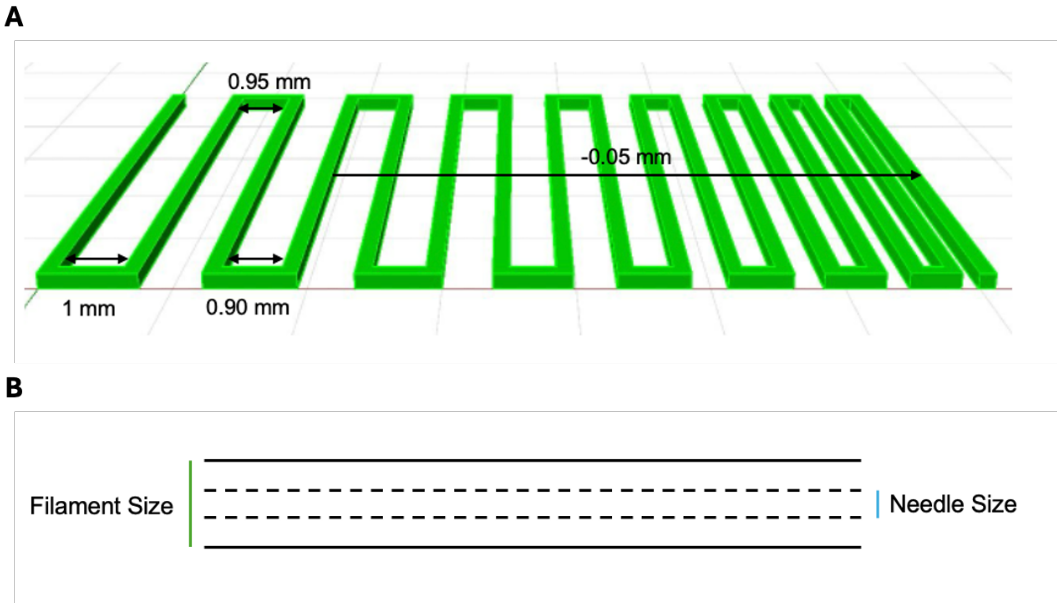


**Supplementary Fig. 1.** Schematic describing the filament fusion test. **(A)** The CAD file measurements and **(B)** an illustration of the needle diameter versus the actual spread of the filament.

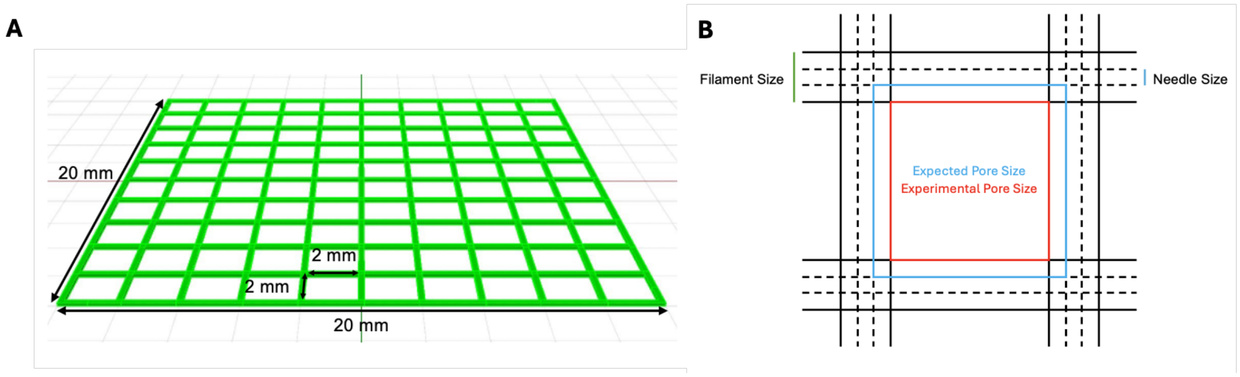


**Supplementary Fig. 2.** Schematic describing the porosity test. **(A)** The CAD file measurements and **(B)** an illustration of the expected pore size versus experimental pore size.

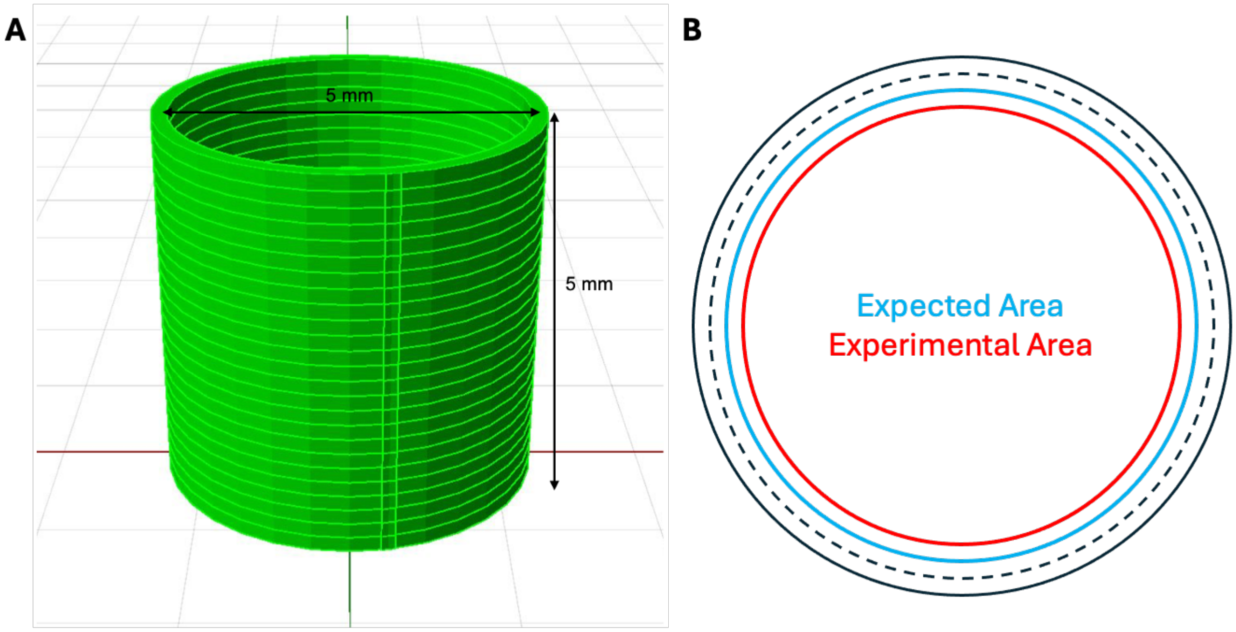


**Supplementary Fig. 3.** Schematic describing the cylinder test. **(A)** The CAD file measurements and **(B)** an illustration of the expected area versus experimental area.

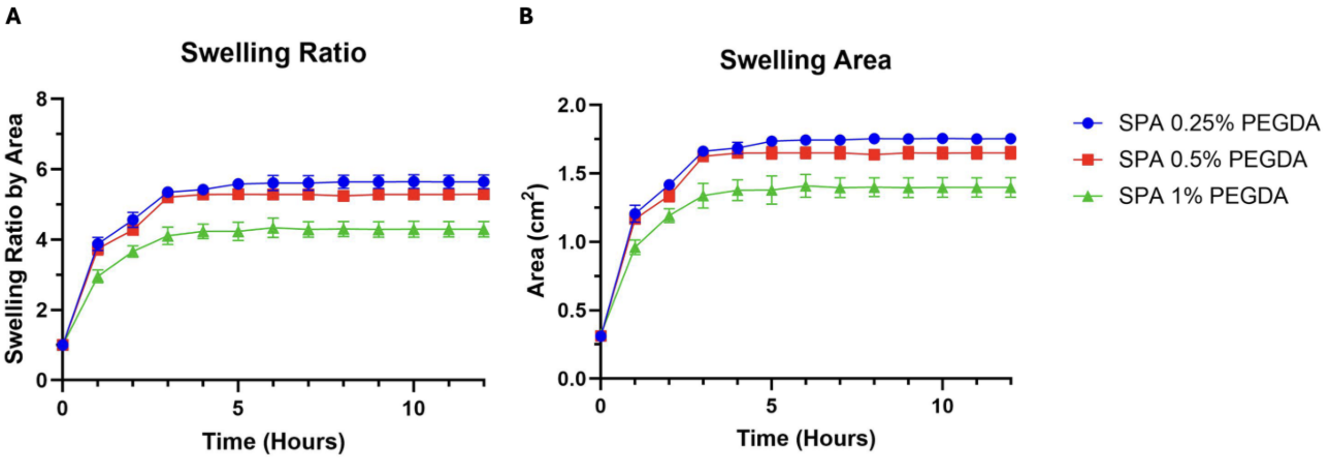


**Supplementary Fig. 4.** Characterization of SPA swelling area over 12 h. **(A)** Swelling ratio by area of SPA with change in PEGDA concentration swelled in DPBS for 12 hours and **(B)** area.

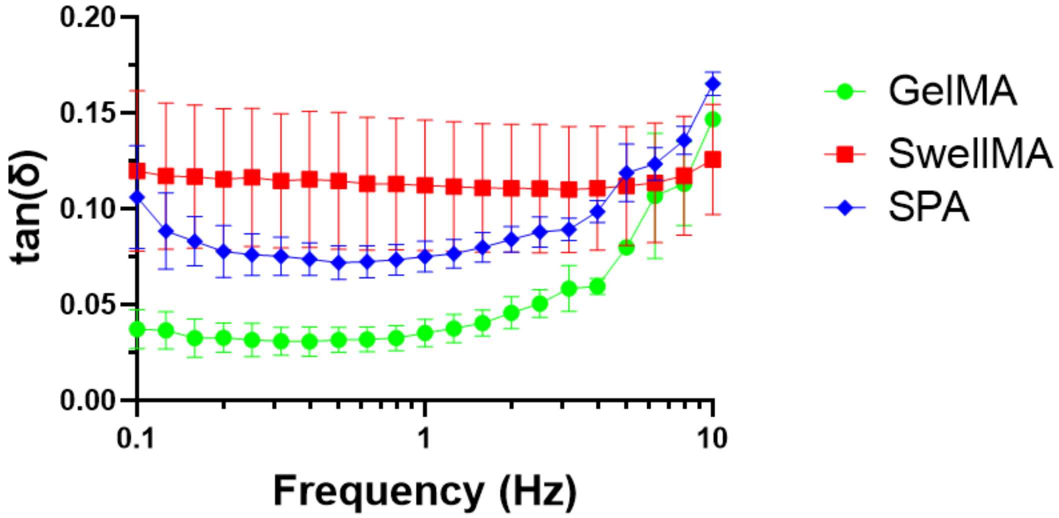


**Supplementary Fig. 5.** Tangent delta (tan (δ)) of GelMA, SwellMA, and SPA throughout the frequency sweep rheological assessment. At 1 Hz, SwellMA exhibits the highest tan(δ) demonstrating its viscoelastic behavior. N = 3, data shown as mean ± SD.

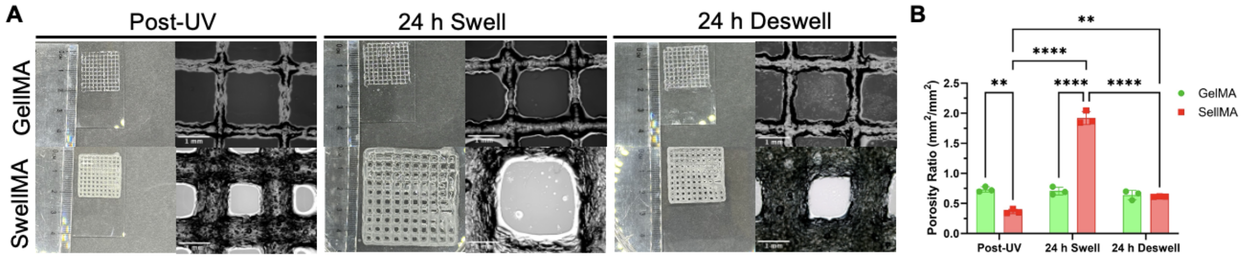


**Supplementary Fig. 6.** Porosity print test of SwellMA and GelMA. **(A)** Macroscopic and microscopic images of the porosity print test for GelMA and SwellMA with 24 h swell and 24 h deswell. **(B)** Quantification of the porosity ratio. **** Denotes significance ($p < 0.0001$), ** ($p < 0.01$), n = 3, data shown as mean ± SD.

**Supplementary Video 1 (.mp4 file).**

**Supplementary Video 2 (.mp4 file).**

**Supplementary Video 3 (.mp4 file).**

**Supplementary Video 4 (.mp4 file).**
